# A Poisson distribution-based general model of cancer rates and a cancer risk-dependent theory of aging

**DOI:** 10.1101/2023.03.08.531809

**Authors:** Wenbo Yu, Tessa Gargett

## Abstract

This article presents a formula for modeling the lifetime incidence of cancer in humans. The formula utilizes a Poisson distribution-based “np” model to predict cancer incidence, with “n” representing the effective number of cell turnover and “p” representing the probability of single-cell transformation. The model accurately predicts the observed incidence of cancer in humans when cell turnover reduction is taken into account. The model also suggests that cancer development is ultimately inevitable. The article proposes a theory of aging based on this concept, called the “np” theory (Nuts Poisoned). According to this theory, an organism maintains its order by balancing cellular entropy through continuous proliferation. However, cellular information entropy increases irreversibly over time, restricting the total number of cells an organism can generate throughout its lifetime. When cell division slows down and fails to compensate for the entropy increase of the system, aging occurs. Essentially, aging is the phenomenon of running out of predetermined cell resources. Different species have evolved separate strategies to utilize their limited cell resources throughout their life cycle.

## 1. Introduction

It has been theorized since the early 1900s that cancer arises from genetic mutations in cells[1–3]. These pioneering works formed the basis of the modern clonal selection theory, which proposes that cancer develops from a single-cell event triggered by a sequence of mutations that transform normal cells into malignant cells[4]. The rate of mutation accumulation is constant throughout the lifespan, which was hypothesized by early theorists and has been further supported by recent evidence[4–6]. A mathematical model of cancer rates, based on the six powers of “t,” was proposed[7], and since then, several variations of this model have been suggested to apply to general or specific cancers [8,9]. All these models utilize the Power function.

Here it is proposed that given that cancer initiation is basically a discrete event, it may be possible to model cancer incidence using a discrete probability distribution, such as the Poisson distribution. There are two levels of discrete events involved in cancerization. At the first level, cancer arises from a single cellular event among the multicellular host. Therefore, the probability of cancer incidence in an individual host can be modeled using the Poisson distribution. At the second level, cumulative genetic mutations are required inside the transformed cell for the transformation to occur. Hence, the probability of a single cell’s transformation can be modeled using the cumulative Poisson distribution function.

This study used a Poisson function model, named as the “np” model, to simulate cancer incidence across a human lifespan. The “n” value represents the effective number of cell turnovers, which was quantified in a recent study [10]. The “p” value represents the probability of a single cell undergoing transformation. When not accounting for the decrease in cell turnover as we age, the model predicted a higher incidence of cancer in individuals over the age of 40 compared to actual observed data. However, by adjusting the cell turnover number, the model accurately matched the observed data. This finding led to the hypothesis that a reduction in cell turnover has evolved to promote longevity. As a result, the study proposed an “np” theory of aging.

The theory of aging can be divided into two main categories: the “programmed” theory and the “wear and tear” theory. The programmed theory proposes that the aging of a species is genetically programmed to adapt its lifespan to its life history within the context of evolution[11,12]. The existence of telomeres provides the best micro-evidence for the programmed theory[13]. On the other hand, the “wear and tear” theory suggests that systems wear out at genetic, cellular, or tissue levels, resulting in aging. There are several sub-theories within this theory, including the somatic mutation theory, which suggests that aging is caused by the gradual accumulation of mutated cells with decreased function[14]. At the non-genetic level, there are various others, including: cross-link theory[15], auto immune theory[16], Glycation theory[17], Oxidative damage theory[18], and molecular inflammatory theory[19]. These theories focus on the micro-mechanism or micro-phenomenon of aging rather than the explanation of the fundamental essence of aging, viz., why aging is inevitable. The disposable soma theory of aging attempts to bridge the gap between the “programmed” theory and the “wear and tear” theory[20]. It suggests that as cells experience increasing wear and tear, maintaining the organism becomes increasingly expensive. Therefore, as a result of natural selection, the organism is eventually abandoned by shutting down cellular maintenance.

Although each theory above explains one or more aspects of aging, none of them can fully explain all the phenomena of aging. In this study, the “np” theory of aging postulates that the risk of cancer is the ultimate restriction to lifespan and uses this perspective to unite most preceding theories of aging.

## 2. Materials and Methods

### 2.1. Images and Graphs

Figure 1A and Figure 2 were plotted by Biorender. Figure 1B was graphed using the Desmos Graphing Calculator (www.desmos.com/calculator).

**Figure 1.**
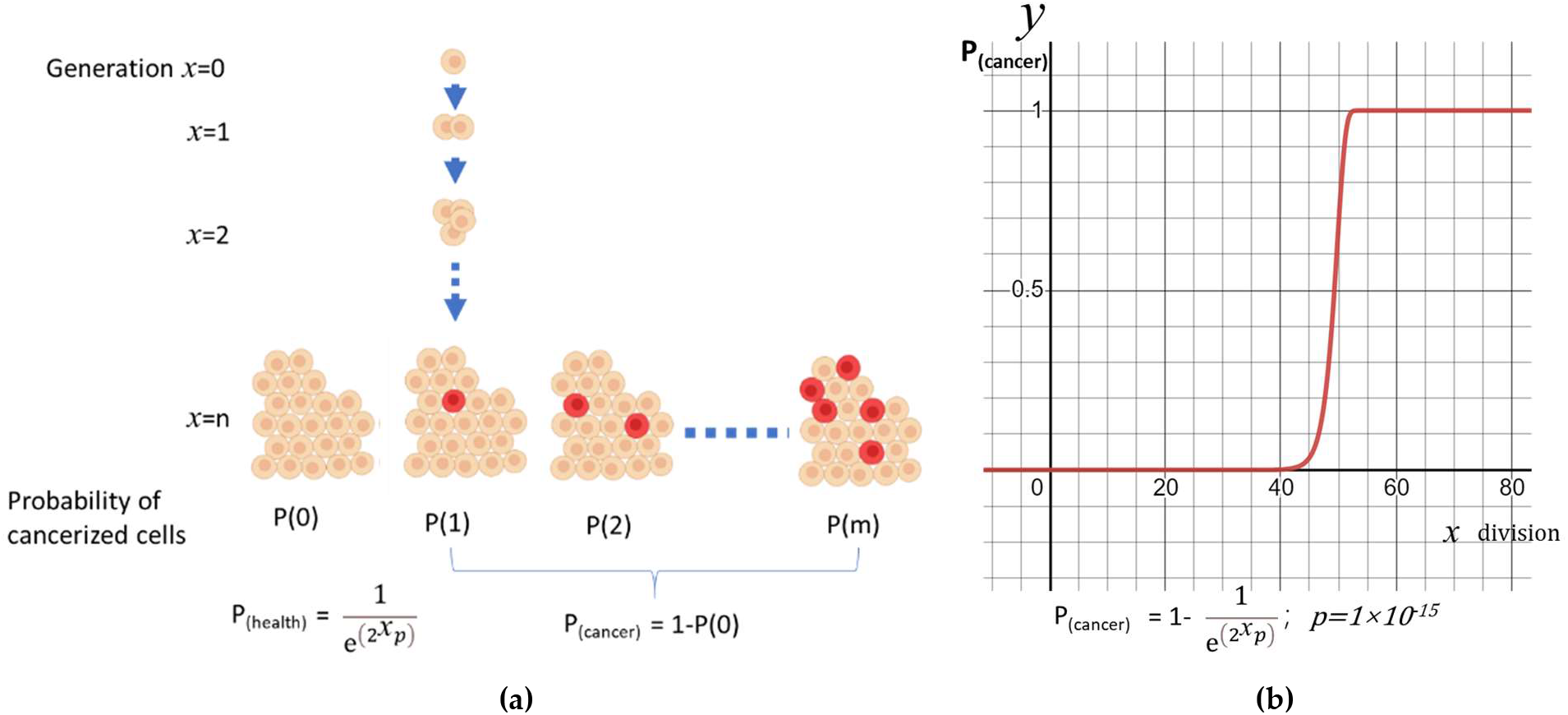
A simple model of cancerization: (**a**) The model of exponentially expanding cell aggregates; (**b**) Probability of cancerization (y) vs division times(x).

**Figure 2.**
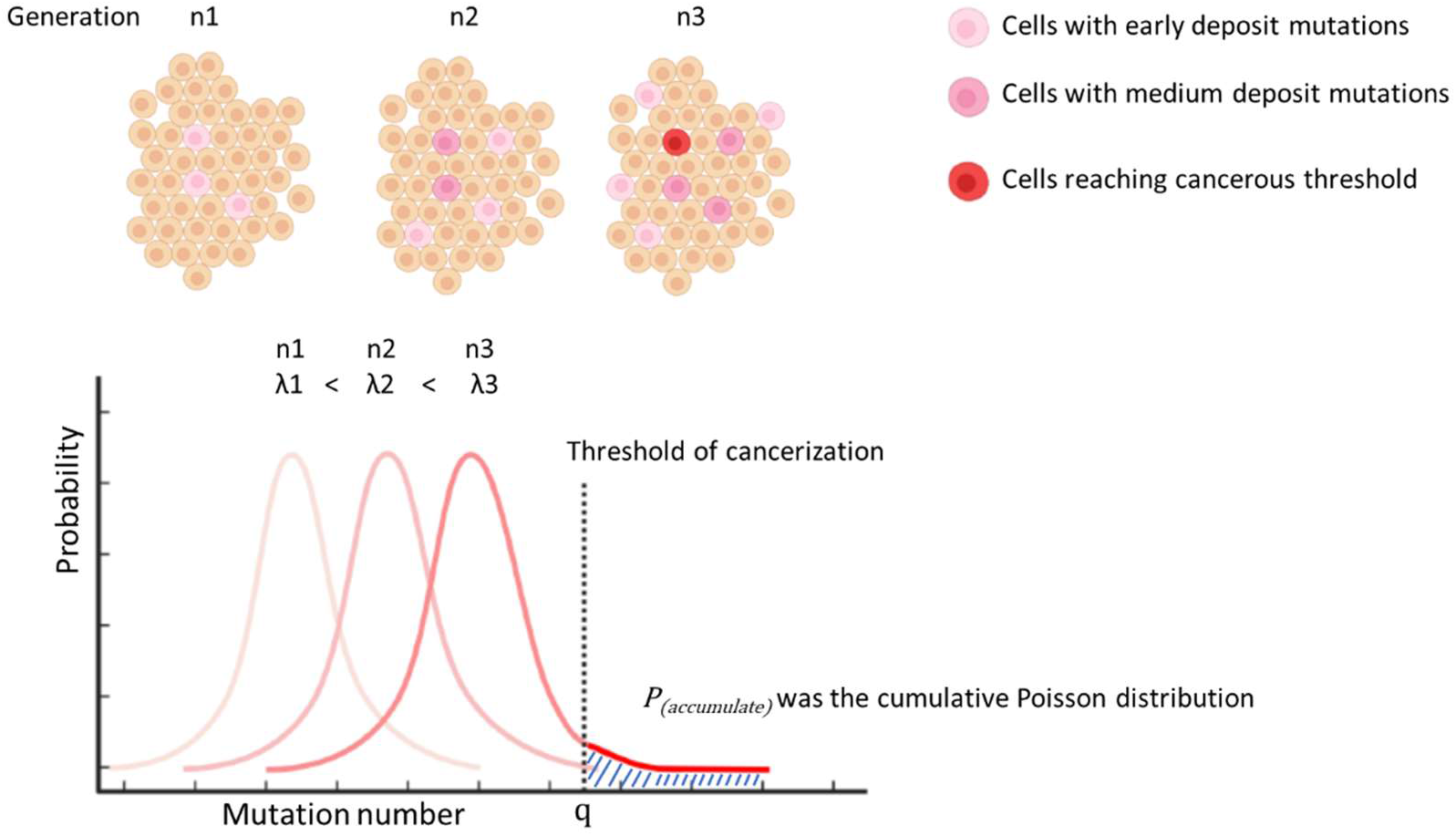
Illustration of modelling P_(accumulate)_ by cumulative Poisson distribution.

### 2.2. Calculation

The Keisan online calculator (https://keisan.casio.com/exec/system/1180573179) was used to calculate the cumulative value of the Poisson distribution p_(a)_ for Table 1 and the supplementary Excel file. The coefficient of determination was calculated as R^2^ = 1-(RSS/TSS). RSS was the sum of squares of residuals, while TSS was the sum of squares of theoretical incidence. 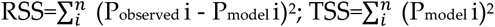.

**Table 1.** The calculation of “np” model.

| Age group | Hypothetic data |  |  |  | Intermediate results |  |  | Final results |  |  | Observed data |
| --- | --- | --- | --- | --- | --- | --- | --- | --- | --- | --- | --- |
| | Generation ($\lambda_{pa}^*$) | n (turnover/year) | $p_{(c)}$ | $p_{(a)}$ | $p=p_{(c)}+p_{(a)}$ | $\lambda=np^*$ | $e^\lambda$ | $P(0)$ | $P(\text{cancer})/\text{year}$ | $P(\text{cancer})/5 \text{ years } (\%)$ | Cancerstats $P(\text{cancer})/5 \text{ years } (\%)$ |
| 0 ~ 5 | 45 | 4.2E+13 | 2.38E-18 | 1.18E-19 | 2.50E-18 | 0.000104939 | 1.000105 | 0.999895 | 0.000105 | 0.0525 | 0.1028 |
| ~ 10 | 46 | 4.2E+13 | 2.38E-18 | 4.69E-19 | 2.85E-18 | 0.000119685 | 1.00012 | 0.99988 | 0.000120 | 0.0598 | 0.0553 |
| ~ 15 | 47 | 4.2E+13 | 2.38E-18 | 2.18E-18 | 4.56E-18 | 0.000191529 | 1.000192 | 0.999808 | 0.000192 | 0.0957 | 0.0633 |
| ~ 20 | 47.5 | 4.2E+13 | 2.38E-18 | 3.12E-18 | 5.50E-18 | 0.000231103 | 1.000231 | 0.999769 | 0.000231 | 0.1155 | 0.1023 |
| ~ 25 | 48 | 4.2E+13 | 2.38E-18 | 6.48E-18 | 8.86E-18 | 0.00037227 | 1.000372 | 0.999628 | 0.000372 | 0.1860 | 0.1643 |
| ~ 30 | 48.5 | 4.2E+13 | 2.38E-18 | 1.33E-17 | 1.57E-17 | 0.000658251 | 1.000658 | 0.999342 | 0.000658 | 0.3286 | 0.3003 |
| ~ 35 | 49 | 4.2E+13 | 2.38E-18 | 2.69E-17 | 2.93E-17 | 0.001230357 | 1.001231 | 0.99877 | 0.001230 | 0.6133 | 0.4533 |
| ~ 40 | 49.5 | 3.15E+13 | 2.38E-18 | 5.38E-17 | 5.62E-17 | 0.001770624 | 1.001772 | 0.998231 | 0.001769 | 0.8814 | 0.6380 |
| ~ 45 | 50 | 2.36E+13 | 2.38E-18 | 1.06E-16 | 1.09E-16 | 0.002569392 | 1.002573 | 0.997434 | 0.002566 | 1.2765 | 0.9550 |
| ~ 50 | 50.5 | 1.77E+13 | 2.38E-18 | 2.08E-16 | 2.10E-16 | 0.003723329 | 1.00373 | 0.996284 | 0.003716 | 1.8444 | 1.5588 |
| ~ 55 | 51 | 1.33E+13 | 2.38E-18 | 4.01E-16 | 4.03E-16 | 0.005361639 | 1.005376 | 0.994653 | 0.005347 | 2.6452 | 2.3953 |
| ~ 60 | 51.5 | 9.97E+12 | 2.38E-18 | 7.66E-16 | 7.68E-16 | 0.00765416 | 1.007684 | 0.992375 | 0.007625 | 3.7548 | 3.5565 |
| ~ 65 | 52 | 7.48E+12 | 2.38E-18 | 1.45E-15 | 1.45E-15 | 0.010820778 | 1.01088 | 0.989238 | 0.010762 | 5.2666 | 5.3138 |
| ~ 70 | 52.5 | 5.61E+12 | 2.38E-18 | 2.70E-15 | 2.70E-15 | 0.015142111 | 1.015257 | 0.984972 | 0.015028 | 7.2915 | 7.5760 |
| ~ 75 | 53 | 4.2E+12 | 2.38E-18 | 4.99E-15 | 4.99E-15 | 0.020971326 | 1.021193 | 0.979247 | 0.020753 | 9.9546 | 9.5098 |
| ~ 80 | 53.5 | 2.73E+12 | 2.38E-18 | 9.11E-15 | 9.12E-15 | 0.024913911 | 1.025227 | 0.975394 | 0.024606 | 11.7123 | 11.8208 |
| ~ 85 | 54 | 1.78E+12 | 2.38E-18 | 1.65E-14 | 1.65E-14 | 0.029297383 | 1.029731 | 0.971128 | 0.028872 | 13.6263 | 13.0510 |
| ~ 90 | 54.5 | 1.07E+12 | 2.38E-18 | 2.95E-14 | 2.95E-14 | 0.031483795 | 1.031985 | 0.969007 | 0.030993 | 14.5654 | 14.2038 |
| ~ 95 | 55 | 5.33E+11 | 2.38E-18 | 5.24E-14 | 5.24E-14 | 0.027916904 | 1.02831 | 0.972469 | 0.027531 | 13.0280 | 13.3100 |

\* $\lambda_{pa}$ was the parameter used in formula (4) to calculate $p_{(a)}$ , $\lambda=np$ was used to calculate the final probability. They are different.

The calculation methods for Table 1: In the Poisson distribution calculator, percentile x=118, mean λ=λ_Pa_ (data from Table 1). p_(a)_ of specific generation was calculated from the difference of neighboured “upper cumulative Q”. For the “0-5” group, put “percentile x” =118 (q), which is the constant threshold. Generation 45 is the “mean λ”. 1.18E-19 is output as upper cumulative Q_(45)_. For next group (5-10), mean λ of 46 is used to get Q_(46)_ = 5.86E-19. The cancerization probability in each age group is (Q_n+1_ - Q_n_)/(1 - Q_n_). Since Q is very small, (Q_n+1_ - Q_n_)/(1 - Q_n_) ≈ Q_n+1_ – Q_n_ = 5.86E-19 - 1.18E-19 = 4.69E-19, which is the p(a) for “5-10” group, so forth, to calculate p_(a)_ for every group. Put “p_(a)_”, “p_(c)_” and “n” into formula (5) to get P(0). P_(cancer)_ /year= 1-P(0). P_(cancer)_/5 years (%) = [1-(1-P_(cancer)_ /year)^5^]× 100.

## 3. Results

### 3.1. A simple model

In a simple model of exponentially growing cell aggregates, the number of cells doubles with each generation “x”. In each division, there is a probability “p” that each cell may experience a cancerous mutation. The probability of “m” cells simutaneously undergoing cancerization out of all the cells follows a Poisson Distribution (Fig. 1a):

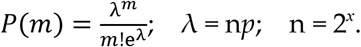

In the healthy group, m=0. So,

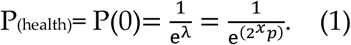

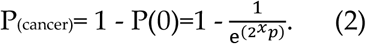

The function will produce an S curve for P(cancer). If we set p=1×10^-15^, the curve will jump to 1 around the 50th generation of division (Fig. 1b). This model suggests that every cellular organism will eventually develop cancer. The likelihood of cancer increases as generations proliferate, with a more rapid increase occurring after a certain age.

### 3.2. An adapted model

Cancer incidence cannot be simply modeled using the formula above because multicellular organisms are not simple cell aggregates that proliferate exponentially without limit, and the “p” value of cancerous mutation is more complex than a constant. We assume that only cells entering the proliferation cycle have the potential for genetic mutation[21]. Hence, “n” equals the cell turnover number during a certain period, which is not constant but rather a function of age “t”, which is corelated to cell generation. “p” is also a function of “t”. The formula is now expressed as follows:

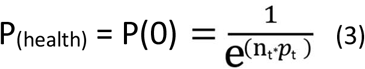

Based on a recent study, the average daily turnover rate of cells in a standard reference person was 0.33 trillion. Of these cells, 65% were red blood cells that lack a nucleus[10], resulting in a turnover rate of cells with active DNA replication of 0.116 trillion per day. For the purposes of this study, a yearly turnover rate of 42 trillion cells will be used for calculations (Table 1). In this model, the parameter “p” is split into two terms: p_constant_ (p_c_) and p_accumulate_ (p_a_). “p_c_” represents the background probability of a single cell becoming cancerous with each division, while “p_a_” represents the probability of cancerization from a cell that has accumulated mutations over multiple divisions. “p_a_” is a function with division generations and is determined based on a raining beads model. In this model, cells are envisioned as infinite bowls into which mutations rain down like beads with each replication. Once the number of mutations exceeds a certain threshold in a bowl, the cell becomes cancerous. The number of mutations in each bowl follows a Poisson distribution (Fig. 2). The probability of exceeding the threshold “q” is calculated as the cumulative Poisson distribution in formula (4):

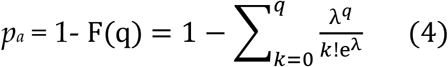

“λ” represents the mean of accumulated mutations per cell. “q” represents the threshold at which a cell becomes cancerous (q > λ). Multiple studies have suggested that somatic mutations increase linearly over the course of an individual’s life [5,6]. Thus, it is reasonable to assume that with each round of replication, the number of mutations also increases proportionally, resulting in “p_a_” increasing as a function of cell division generation or time (Fig. 2). From formula (3), a new formula can be derived as follows:

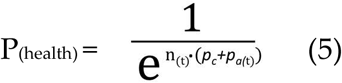

Here we set p_a(t)_ as an internal parameter that does not need to have a specific biological meaning. This internal parameter p_a(t)_ is used to demonstrate that the overall cancer incidence follows the cumulative Poisson distribution.

### 3.3. Fitting the model to observed cancer incidence

We retrieved the data of the average number of New Cases Per Year and Age-Specific Incidence Rates per 100,000 Population in UK (cancerstats)[22]. We used these data to fit our proposed model formula (5).

#### n

Since the turnover number “n” was obtained from the reference Man aged between 20–30 years, we will apply “n” to the group up to age 35 (Table 1). We have no data on cell turnover in children. Since “p” is very low in the early stage of life, the impact of “n” is limited. Furthermore, considering the higher metabolism status and smaller body mass of young children, we will keep “n” the same value before age 35.

#### p_c_

In the early stages of life, “p_a_” was insignificant, and we estimated P_(cancer)_ as 0.05% per year based on Cancerstats data. Based on formula (5) (Supplementary Excel), “p_c_” can be deduced as 2.38E-18.

#### p_a_

As previously discussed, the exact biological meaning of “p_a_” cannot be provided at this stage. It is an internal parameter that reflects the increasing probability of cancer incidence, based on the assumption that, on average, each generation of cell division will randomly deposit equal amounts of cancer-related mutations following the Poisson distribution[23,24]. “λ_Pa_” represents the mean number of mutations for each cell, and cells will become cancerous when the number of deposited mutations reaches the threshold “q”. To calculate “p_a_” using formula (4), we set “λ” equal to the cells’ generation (Table 1: λ_Pa_) and tried different thresholds “q” until the model best matched real cancer rates (with the highest R^2^). Ultimately, we set “q = 118”, which means that a cell requires 118 mutations on average to become cancerous, assuming it receives one mutation from each division (Fig. 3A). Determining the generation of cells at different ages presents a challenge because cells from various tissues may have different developmental histories and, as a result, different generation numbers. Additionally, differentiated and stem cells may have distinct division cycles. However, we provided an average estimate of generations to assist in building the model and prove that cancer incidence follows our mathematical hypothesis. From the fertilized egg to the newborn infant, cells proliferate exponentially, and the newborn has a total of two trillion cells [25], meaning it has undergone 41 generations of divisions (supplementary excel). For the first five years of life, it requires at least another four generations, and we set λ_Pa_ = 45 for this age group. For the 5-10 and 10-15 age groups, we set one generation for each stage. Above this age, we set 0.5 generation for each stage until it reached the Hayflick limitation of 55[26].

**Figure 3.**
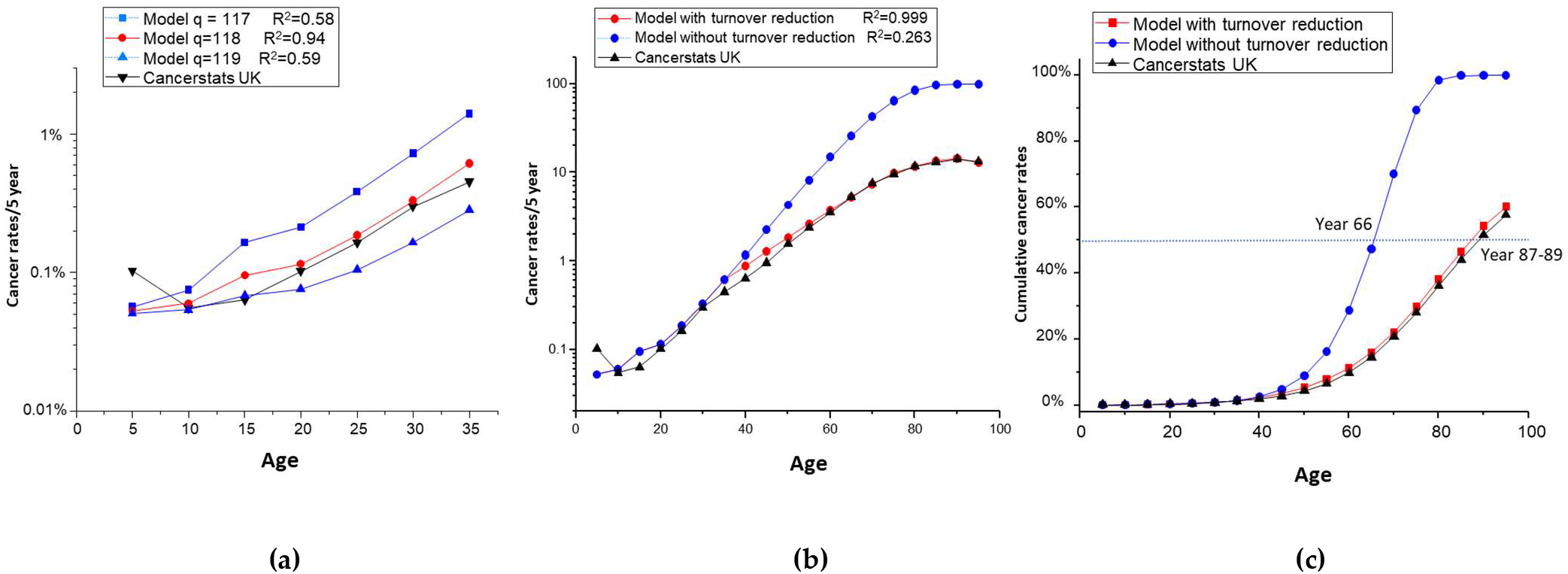
Match model predicted data with real data: (**a**) Different “q” were tested for the formula using groups year 0-35. q=118 was chosen for the final model; (**b**) All life span model of 5 years’ cancer incidence with or without considering cell turnover reduction; (**c**) All life span model of cumulative cancer incidence with or without considering cell turnover reduction.

By setting n, p_c_, p_a_, and using formulas (4) and (5), we can model the five-year cancer incidence (Fig 3A and Table 1). We used q=118 for further analysis.

### 3.4. Final adaption of the model to account for reduced cell turnover

The predicted incidence of cancer exceeded the observed data beyond the age of 35 (Fig 3b). This occurred because Formula (5) cannot always use the same “n.” As people age, cell division and turnover rates decrease[5]. As no real data on cell turnover in aging people are available, we determined the turnover decrease rate by assuming the validity of our model. We found a 25% decrease per five years in the 35-75 age group, a 35% decrease per five years in the 75-85 age group, a 40% decrease per five years in the 85-90 age group, and a 50% decrease per five years in the group aged over 90 years (Table 1: n turnover/year). The model accurately fits the observed data since the reduction was reversely deduced (Fig 3b). Therefore, it is feasible to use a general theory-based model to match cancer incidence. This model authentically reflects the decreased cancer incidence in the very aged group[5]. Several studies have reported that cancer rates exhibit exponential growth by six powers of “t” between ages 20 to 75 [2,3,7]. Fisher and Hollomon’s pioneering study of stomach cancer found that ΔLog(p)/ΔLog(age) has a slope of 5.7 between the ages of 20-75[2]. It is worth noting that the “np” model, without considering cell turnover reduction, also yielded a straight line with a slope of 5.67 from Group 25 to Group 75, which precisely matches Fisher’s case (Supplementary Excel). This implies that stomach tissue may not experience an apparent reduction in cell turnover during this age period.

### 3.5. A theory of aging based on the cancer model

If we convert the cancer incidence shown in Fig 3b into cumulative incidence, we get Fig 3c. From this figure, we can see that reduced cell turnover actually offers a greater advantage in terms of survival. The model indicates that without cell turnover reduction, humans would reach a 50% cancerization rate at age 66, but with cell turnover reduction, the 50% cancerization rate is delayed by two decades to age 87-89 (Fig 3c). This gives us a hint of the ultimate cause of aging, which is based on the unavoidable increase of cancer risk.

Here, we propose an “np” theory of aging. Cells are highly ordered systems, and to maintain cell fitness (youth), the order needs to be maintained, which can be described as an issue of entropy balance[27]. A cell always gains positive entropy, which needs to be reconciled to defy the second law of thermodynamics. Three levels of entropy are postulated here: (1) metabolic entropy; (2) structural entropy; and (3) information entropy. (1) For any living cell, metabolism is the function to maintain energy/matter intake and output. The entropy at this level is balanced biochemically. (2) With time, the microstructure of the cell or cellular organelles experience “wear and tear”. The generation of new cells through division is the final resort to fix this “wear and tear” and reduce structural entropy. (3) However, the irreversible random changes accumulated in the genetic material that cannot be fixed will be passed to the progeny cell. The accumulated information entropy will ultimately succumb to the second law of thermodynamics. The increase in information entropy finally destabilizes the regulation of the cell and leads to unregulated proliferation, resulting in cancer[28,29]. From another perspective, we can categorize cellular information into two arms: pro-proliferation and pro-regulation. The genetic mutations randomly impact either arm, but only the disruption to the pro-regulation arm will be selected for. With the increase of information entropy, the highly regulated eukaryotic cells will return to a more primitive prokaryotic-like status[28]. The beast of proliferation will be released. The postulate of the information entropy predicts that any multicellular system will eventually develop cancer. As a result, the total number of cells that can be usefully generated from a single zygote is finite. To minimize the risk of cancer, at the later stage of a species’ lifespan, cell turnover is reduced or stopped. The negative entropy introduced into the cells cannot balance the positive entropy produced by the system, leading to increased disorder in cellular structure and metabolism. When this happens, the entropy of the whole system increases, the fitness of the organism decreases, and aging occurs.

This theory of aging predicts the ultimate number of cells a given individual can use is “N”. “N” is restricted by “p”. The predetermined number “N” can be plotted as an enclosed area on the “n” and “t” graph (Fig 4a). For the same reason, the quality of reproductive cells is also restricted by the same law[30]. Hence, any species has a certain period of reproduction. Species will develop different ways to use this cell resource strategically, which constitutes the basis of an organism’s lifespan and aging process. We list three models of species with three typical lifespans for their survival strategies.

**Figure 4.**
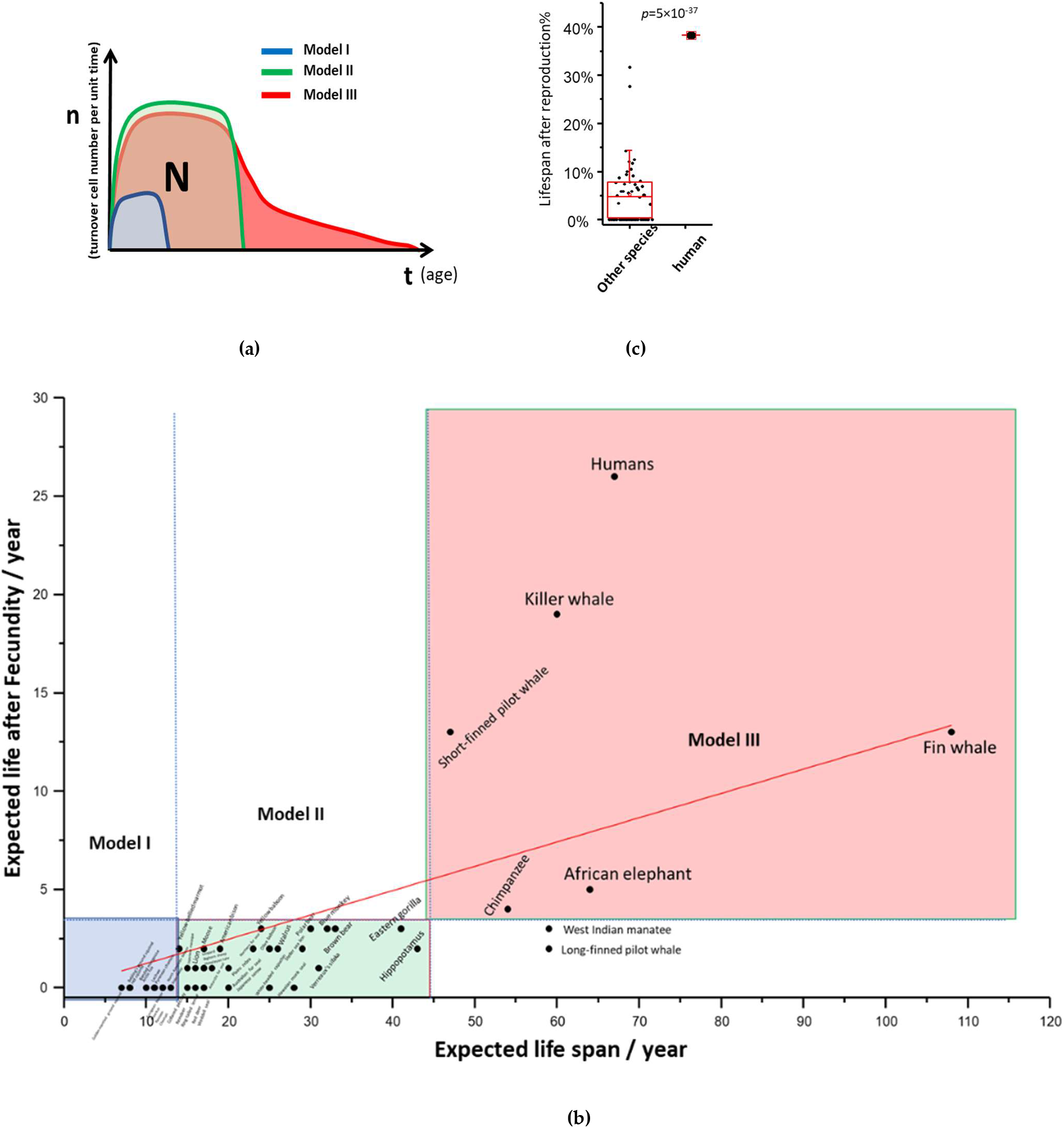
“np” theory of aging among different species: (**a**) Theoretical “nt” plot of model I, II, III species; (**b**) Post reproduction life vs expected life span of 51 mammal species; (**c**) The percentage of post reproduction life to whole life: human against the other mammals. T test was used to calculate statistical significance.

Model I: Species with short lifespan and short post-fecundity life. Low fitness was not acceptable for these species. Model I species have a very short half-life of survival in the natural environment, so there is not much evolutionary pressure for longevity. Their lifespan is compatible with their survival probability. The “nt” curve has a small area on the plot. The model species are rodents.

Model II: Species with medium to long lifespan and short post-fecundity life. Low fitness was not acceptable. If the species adapt to a strategy where longevity is favored, they are allowed to have more “N,” which enlarges the enclosed area on the “nt” plot. This process can continue under evolutionary pressure until the advantage of longevity is canceled off by the cancer risk. These species are stronger and have a higher chance of survival for a longer period, so evolution gives them more predetermined cells in their lifespan. However, lifespan is still restricted by the cancer risk. Eventually, the organism will shut down cell proliferation quickly and exit from survival competition. The model species are large carnivores.

For Models I and II species, after the reproductive period, the organism becomes aged and dies quickly. They cannot evolve the strategy of reduced cell turnover because reduced turnover means reduced fitness, and fitness is crucial for their survival. Their lifespan matches the disposable soma theory[20].

Model III: Species with long lifespan and a long post-fecundity life. Low fitness is acceptable. Few species are extremely favored by longevity. This strategy may be evolutionarily favored by the “grandma effect”[31–33], where longevity may provide community benefit. We hypothesize that the “N” reaches an evolutionary limit, but the Model III species develop another strategy for using the available “N” by reducing cell turnover at the cost of lower fitness. This type of species has an elongated senescence period among all species. All of them are social and intelligent species. Surviving with low fitness is possible in the community. This also offers an explanation for the brain weight theory, which found that lifespan was positively related to species’ brain weight[34].

To support this theory, we re-explored the data from Samuel Ellis and Darren P. Croft about the reproduction lifespan and post-reproduction lifespan of 51 mammals[35]. The post-reproduction lifespan vs. total expected lifespan was plotted (Fig 4b). If we divide the species into three groups based on their expected lifespan on the x-axis and two groups based on post-reproduction life on the y-axis, 49 out of 51 species fall into three groups (Table S1). These three groups represent aging strategy models I, II, and III, respectively. We note that humans have the highest post-reproductive lifespan and the highest percentage of post-reproductive time (Fig 4c), suggesting that humans have a unique position in evolution and that longevity is highly favored in this species.

## 4. Discussion

This study describes a model of cancer incidence that give arise to a wider theory of aging. It’s important to note that “p” should not be simply interpreted as the rate of DNA mutation. Instead, it represents the overall likelihood of a cell to escape regulation or suppression and develop into a cancerous colony. The development of cancer is influenced by complex factors, although growing evidence supporting random mutation as the major contributor[36]. These factors eventually converge at the genetic level, which is represented as “p” in the proposed model. The aim of this mathematical model is to demonstrate that there is a unifying law behind these diverse factors that drive the average pace of cancerization.

While many studies on cancer origin focus on stem cells, it’s crucial to note that all transit-amplifying cells can potentially transform into cancerous cells by dedifferentiation[37]. Therefore, the model includes the number of cellular turnovers, denoted as “n,” which should be proportional to the number of cancer progenitors.

When considering the “np” in different tissues, it is important to view an organism as a developing tree, where the branches may not all develop at the same pace. While this study presents the average expression of “np,” in reality, different tissues may have varying “np” values due to developmental asymmetries in cell lineage trees[38]. This presents an opportunity to further adapt the model for tissue-specific cancers such as breast or prostate cancer. Additionally, this could explain the relatively high incidence of some cancers in children. For example, during early development, the nervous system branch undergoes more divisions than other tissues and accumulates a higher “p,” which slows down after adulthood. The early accumulation of mutations in brain tissue can explain the early peak of brain tumor incidence in children. This principle can also apply to explaining the increased risk of lymphoma observed in AIDS patients or the positive relationship between chronic inflammation and cancer[39,40], as these diseases lead to increased cell turnover.

While our discussion has been centered on mammals, this theory may also apply to other vertebrates. For instance, long-lived, cold-blooded animals like turtles or the Greenland shark have a slower “cellular clock” and slower metabolism, which may lead to longevity[41,42].

Over the last decade, DeGregori et al. developed a theory of cancer development based on the fitness of cancer progenitor cells, which was actually an attempt to apply the disposable soma theory in tumorgenesis[43–48]. According to this theory, genetic mutation is not the primary driver of tumor development. Instead, the mutated cells are suppressed by the hosts until the post-reproduction period, when the host relaxes tumor repression. The theory suggests that normal stem cells have a higher fitness in young tissue environments, which makes it difficult for mutant progenitor cells to compete with healthy stem cells. However, as the system ages, the microenvironment changes, and the healthy stem cell loses its competitive advantage. Mutated cells then gain relatively higher fitness than normal stem cells, leading to tumorigenesis. One problem with the theory is the lack of evidence to support the micro-mechanism. There is evidence to support either a gain or loss of fitness in mutant cells, and there could be many mutations with little phenotypic or fitness change[49]. The disagreement here is obvious: the “np” theory postulates that cancer is the ultimate restrictor of lifespan, and aging was a strategy to avoid cancer, while DeGregori’s theory postulates that aging relaxes the soma regulation thereby allowing cancer development.

There may be ways to resolve this argument. If we can identify the molecular clock that regulates a particular tissue, we can slow down the turnover of stem cells in that tissue[50,51]. For example, if we slow down the stem-cell turnover in mouse breast tissue, based on the “np” theory, we would expect the tissue to display signs of aging but maintain genetic youthfulness, which could delay breast cancer by promoting aging. However, if DeGregori’s theory is correct, this practice would have no impact or could even promote cancer, since aged tissue relaxes its control of tumorigenesis (Fig 5).

**Figure 5.**
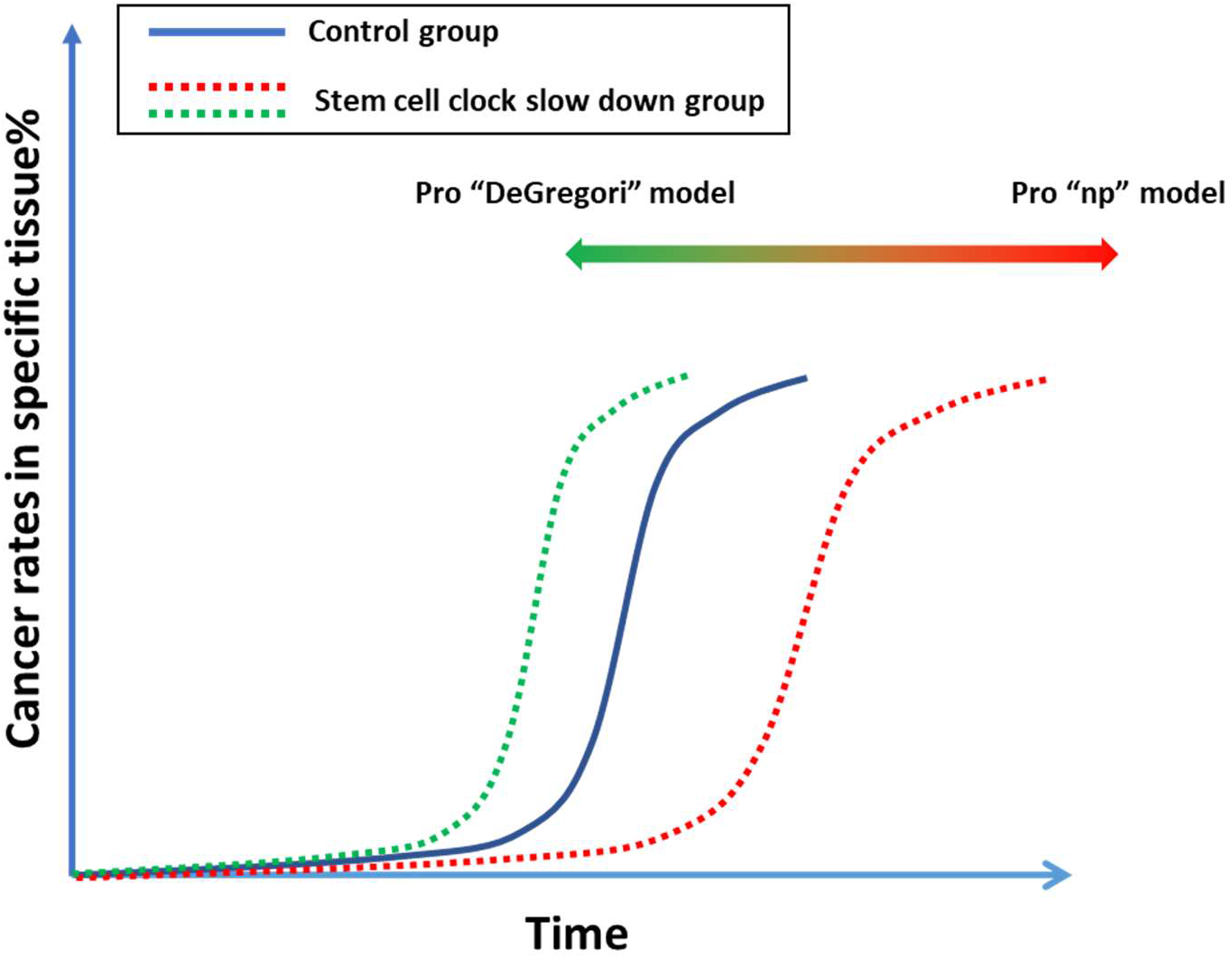
A proposed experiment which can possibly resolve the argument of “np” theory and DeGregori’s theory.

We can use the “Nuts Poisoned (np)” model as a metaphor for the aging theory. Imagine a tree of life that produces “Nuts” (fresh cells with low entropy) that support life. A creature feeds on these nuts, which help maintain its fitness. However, some nuts may be poisoned, and over time, more nuts will get poisoned. To increase the chances of survival, the creature must reduce its nut consumption to minimize the risk of poisoning. However, this reduction in nut consumption causes the creature’s fitness to decline, and it begins to age. Eventually, the creature must abandon the tree of life because it has become too poisonous.

We propose that aging is a manifestation of entropy increase. The accumulation of system entropy can be observed as aging[52]. A study of bacterial aging has shown that cells can balance their entropy by proliferating[53]. However, the mechanism of how proliferation can restore negative entropy is not fully understood. Some studies have suggested that division can reduce entropy by altering the cells’ surface-to-volume ratio or through compartmentalization[54,55]. Our very existence from the first cell on earth demonstrates that cells can renew themselves indefinitely. Information entropy measures the quality of genetic material, which cannot be perfectly maintained forever. Therefore, the ultimate limitation on life is information entropy. The only way to overcome this limitation is through single colony selection, and the process of reproduction is just such a form of single colony selection. Natural elimination of imperfect seeds maintains the stability of information entropy from generation to generation.

Many scientists believe that biological systems have the inherent ability to repair damage and replace defective cells, which suggests that they are not necessarily destined to die[12]. However, the accumulation of genetic mutations is an inevitable process that affects every living organism, leading to mortality. Although stem cell therapies hold promise, they have also been associated with the side effects of tumorigenesis, which can be explained by our theory[56,57]. As a Model III species, humans have almost reached the ceiling of lifespan, implying that modern antiaging practices such as calorie restriction[58], anti-oxidant[59], Rapamycin or Sirtuins treatment, will not break the ceiling to extend life beyond this limit[60,61]. Cancer incidence is not significantly different between small, short-lived animals and large, long-lived animals, as all species have evolved to adapt their lifespan to their available resources. Thus, the “np” theory offers an explanation for Peto’s Paradox[62].

Despite the challenges, there is still hope. If the “np” theory is correct, it could provide new insights into cancer prevention and human longevity. According to the formula, the strategy would be to reduce “p” and “n”. To prevent specific cancers, one approach could be to slow down the stem cell clock in the tissue (low “n”). Alternatively, if low fitness is unacceptable, we could replace the tissue with fresh stem cells. We should develop techniques for identifying stem cell colonies ex vivo to ensure that they have the perfect genome (low “p”). Similarly, we could develop anti-aging technologies based on the same principle. However, ethical issues must be carefully considered.

## 5. Conclusions

In this study, we formulate the first general model for cancer incidence across all lifespans based on Poisson distribution. We propose a new theory of aging, known as the “Nuts Poisoned” theory, which aims to address gaps in existing aging theories. Aging is fundenmentaly tangled with the inherent risk of cancer. We discuss various scientific observations in the context of this theory and suggest new avenues for cancer prevention and anti-aging strategies. Currently, this theory is applied only to mammals, but it has the potential to be extended to other vertebrates as well.

## Notes

### Competing Interest Statement

The authors have declared no competing interest.

