## Supplemental Table for "A Poisson distribution-based general model of cancer rates and a cancer risk-dependent theory of aging"

### Slide 1
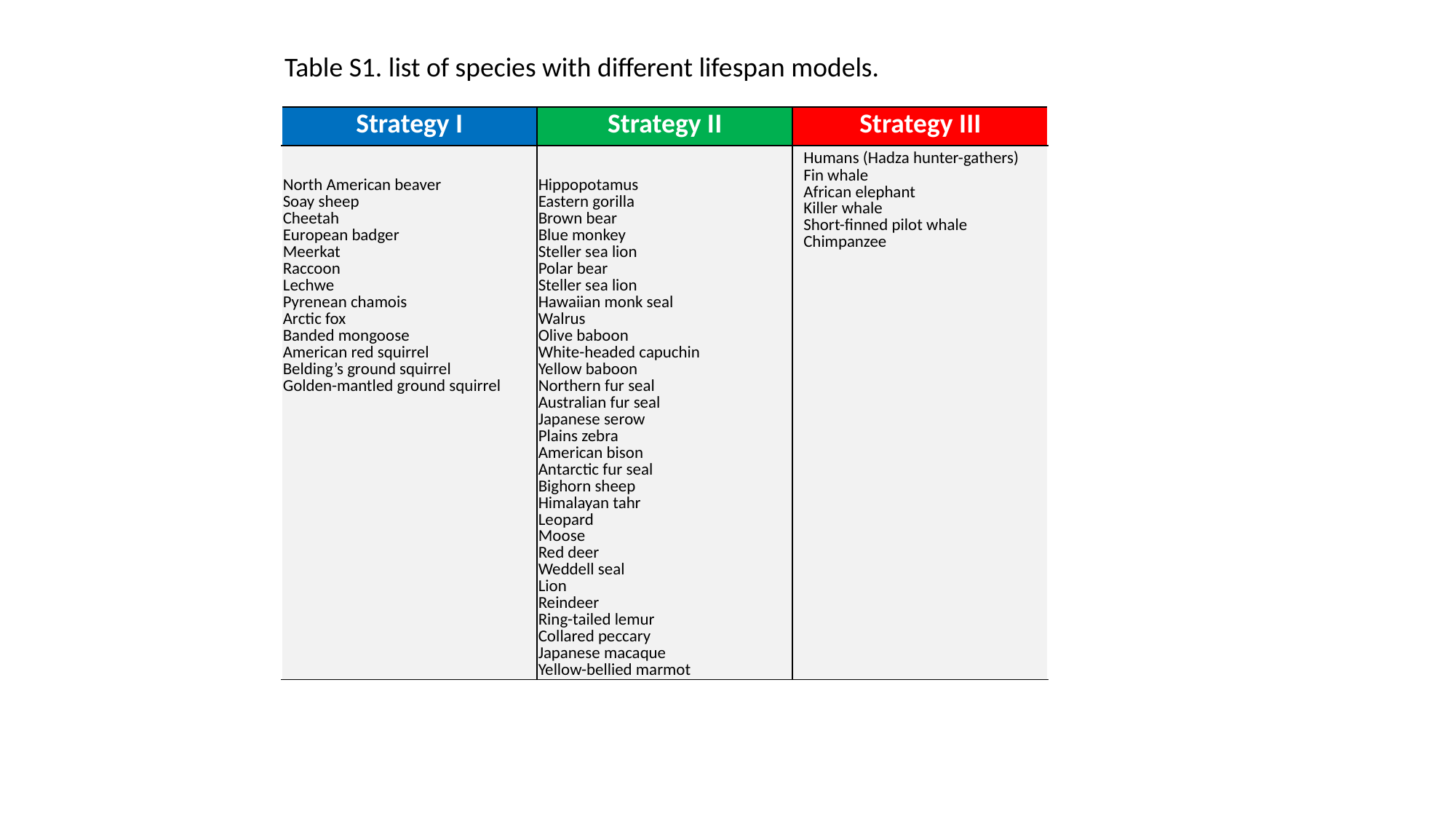

Table S1. list of species with different lifespan models.
| Strategy I | Strategy II | Strategy III |
| --- | --- | --- |
| North American beaver Soay sheep Cheetah European badger Meerkat Raccoon Lechwe Pyrenean chamois Arctic fox Banded mongoose American red squirrel Belding’s ground squirrel Golden-mantled ground squirrel | Hippopotamus Eastern gorilla Brown bear Blue monkey Steller sea lion Polar bear Steller sea lion Hawaiian monk seal Walrus Olive baboon White-headed capuchin Yellow baboon Northern fur seal Australian fur seal Japanese serow Plains zebra American bison Antarctic fur seal Bighorn sheep Himalayan tahr Leopard Moose Red deer Weddell seal Lion Reindeer Ring-tailed lemur Collared peccary Japanese macaque Yellow-bellied marmot | Humans (Hadza hunter-gathers) Fin whale African elephant Killer whale Short-finned pilot whale Chimpanzee |
